# Assessment and comparison of two different methods to extract nucleic acids from individual honey bees

**DOI:** 10.1101/2020.12.21.423857

**Authors:** Rohan Swami, Brooke Ganser, David R. Tarpy, Micheline K. Strand, Hongmei Li-Byarlay

## Abstract

The honey bee is an excellent model system to study behavior ecology, behavioral genetics, and sociogenomics. Nucleic acid based analyses enable a broad scope of research in functional genomics, disease diagnostics, mutant screening, and genetic breeding. Multiple levels of analysis lead to a more comprehensive understanding of the causes of phenotypic variation by integrating genomic variation, transcriptomic profiles, and epigenomic information. One limitation, however, is the sample preparation procedures to obtain high quality DNA and RNA simultaneously, particularly from small amounts of material, such as tissues of individual bees. We demonstrate that it is feasible to perform dual extractions of DNA and RNA from a single individual bee and compare the quality and quantity of the extracted nucleic acids using two different types of methods (column based versus ethanol precipitation). We obtained a higher yield of both DNA and RNA with one of the extraction methods over the other, although the quality of the DNA and RNA was not significantly different between methods. We describe here the first validated method for dual extraction of DNA and RNA specifically from individual honey bees.

## 1. Introduction

Honey bees (*Apis mellifera*) are one of the primary species of social insects used for genomic biology studies. Their caste-specific social behavior, phenotypic plasticity, and genetics are different from other non-social model insect species such as *Drosophila*, and they were one of the first insects to be seqeuenced (Weinstock et al., 2006). Continued interest in honey bee health and the availability of its genome (Elsik et al., 2014) make it a key genomic study organism for the foreseeable future. In addition to studies using the honey bee as a model system to study social evolution, behavior ecology, physiology and genetics, this species is also an excellent model species for behavioral epigenetics (Yan et al., 2015, Yan et al., 2014, Li-Byarlay, 2016) with the first described functional CpG methylation system in insects (Wang et al., 2006). CpG methylation plays an important role in the nutritional control of reproductive status to develop into a queen or a worker bee (Kucharski et al., 2008), as well as the phenotypic plasticity of workers (Herb et al., 2012). Simultaneous studies at the DNA and RNA level revealed that CpG methylation alters gene splicing in honey bees (Li-Byarlay et al., 2013, Foret et al., 2012, Wang & Li-Byarlay, 2015).

Honey bees are also the most important managed pollinator in agriculture, required for the pollination of an annual $15 billion worth of agricultural crops (Bond et al., 2014) and responsible for pollinating a third of human food crops (Gallai et al, 2009). Thus the ongoing annual 30-40% mortality of honey bee colonies (Kulhanek et al., 2017) is a major concern. Several factors are thought to contribute to this mortality, including parasitic mites, pathogens such as bacteria and viruses, pesticide use in agriculture, climate change, poor nutrition, and low genetic diversity (Wang & Li-Byarlay, 2015, Gisder & Genersch, 2017, Simone-Finstrom et al., 2016, Goulson et al., 2015, Wu et al., 2011, Degrandi-Hoffman et al., 2015, Rinderer et al., 2010, Hamiduzzaman et al., 2017). Understanding the molecular biology and genomics of honey bees is crucial to countering the threats to honey bee mortality (Grozinger & Robinson, 2015).

Genome sequencing and transcriptome studies in honey bees have become increasingly common since the completion of the honey bee genome in 2006 (Weinstock et al., 2006). However, the processes of obtaining experimental samples and tissues are not always consistent. Typically, DNA and RNA material are often not obtained from the same specimen but instead are extracted from different pools of specimens. This creates variation that can cause misinterpretation or masking of important data due to variation among individuals (Sultan et al., 2007). Using pooled samples can negatively affect the robustness, resolution, or expressiveness of experiments (de Jong et al., 2010). It is therefore highly beneficial to obtain both genomic and transcriptomic information from the same individual for a truly integrative study of the genome and its transcriptome product.

The extraction of DNA and RNA molecules is the most critical method used in genomic biology (Tan & Yiap, 2009) because any errors at the start will have amplified effects for downstream sequencing procedures and data analysis. Extracting high quality DNA and RNA simultaneously is also central to the studies of gene expression and gene regulation to address phenotypic plasticity and social behavior of honey bees, for example by eQTL mapping (Gilad et al., 2008), as well as to other interesting areas of *Apis mellifera* biology such as the high recombination rate (Rueppell et al., 2016, Wallberg et al., 2015, Beye et al., 2006, Wilfert et al., 2007). Dual extraction is the process of isolating the DNA and RNA from the same sample, eliminating genetic masking from pooled samples, and allowing direct comparisons between the RNA and DNA to be made at the individual level (Triant & Whitehead, 2009). Studying DNA and RNA from the same individual facilitates a more accurate interpretation of the relationship between genotype and phenotype and, therefore, a potentially deeper understanding of the molecular basis of diseases and other complex traits (Reuter et al., 2016). Since dual extraction is not yet a standardized method in honey bees, researchers typically use DNA and RNA from different bee samples to carry out singular extractions. However, there are dual DNA and RNA extraction kits available for purchase on the market today. To our knowledge, these kits have not been systematically compared in honey bees, but they hold great promise and have been used on plant and animal tissues, bacteria, yeast, fungi, algae, viruses, cultured cells, and other insects.

Here, we address two important questions for implementing dual extraction in honey bees. First, we determine whether dual extraction is feasible using honey bee samples from three different life stages (larva, pupa, and adult). Second, we quantify the differences in terms of yield and purity with the different extraction procedures. We demonstrate that the dual extraction technique can be applied to individual honey bees and produce meaningful quantities of DNA and RNA, which can be used for truly integrative genomic analyses at the individual level.

## 2. Materials and Methods

### 2.1. Samples

All honey bee eggs, larvae, pupae, and adults were acquired from hives at the North Carolina State University apiary in Raleigh, North Carolina. Additional adult samples were collected from the Central State University apiary in Wilburforce, Ohio. Five to eight samples were taken for each developmental stage: larval (fifth instar, L5), pupal (brown eye stage), and adult heads (one day old). Before running one dual isolation per sample with each kit, the samples were flash frozen and stored in a −80 °C freezer.

### 2.2. Method 1 for dual extraction

The Bio Basic dual extraction kit was purchased from Bio Basic (BS88203, Toronto, Canada). This extraction kit uses a column technique to isolate DNA and RNA from samples, and we followed the manufacturer’s instructions. The weight of each sample is within 25-50 mg following the manufacturer’s guidelines (Supplemental Table 1). Frozen samples were homogenized and lysed to break open cells. Homogenization was carried out using a plastic micropestle to grind tissues in a 1.5ml micro-centrifuge tube using a cordless motor. After homogenization, lysis buffer was added to the sample and further homogenized with a pellet pestle in order for maximum extraction. During the initial stages of the protocol, DNA and RNA were separated by specifically binding DNA to a column while RNA was in the flow-through supernatant. All samples were centrifuged at 4 °C to ensure binding of DNA or RNA to the spin column inside of the collection tube. Total DNA was obtained after washing the DNA column and elution with buffer. RNA was obtained from DNA flow through after elution wash, received a wash in 70% ethanol, and separately bound to an RNA-specific column, which was then similarly washed and eluted.

### 2.3. Method 2 for dual extraction

The MasterPure dual extraction kit was purchased from Epicentre (MC85200, Wisconsin, USA). The weight of each sample is within the range of 2-5 mg according to the manufacture’s guidelines (Supplemental Table 1). The homogenization and lysation were carried out using the pestle and 1.5ml microcentrifuge tubes. Tissue samples were dipped into liquid nitrogen, homogenized in Proteinase K and cell lysis solution with the use of a pellet pestle, further incubated at 65 °C for 15 minutes, vortexed at 5-minute intervals, and the samples were then stores on ice. Samples received a precipitation reagent and were centrifuged at 4 °C before receiving isopropanol. DNA and RNA were retained in the supernatant, then after a precipitation and wash, total nucleic acid was treated with RNase to obtain total DNA and treated with DNase to obtain total RNA. All samples received two washes with 70% ethanol before being suspended in buffer for analysis and storage. RNA samples also received an RNase inhibitor for protection from degradation while being stored in buffer solution.

### 2.4. Analyzing Yield and Purity

Purity and quantity of the resulting DNA and RNA samples were assessed by photospectrometric analysis using a NanoDrop™ Spectrophotometer (ND-2000, Thermo Scientific, USA). The purity was determined as the 260/280 ratio and the internal Nanodrop™ quantification algorithm was used to determine quantity.

## 3. Results

To compare the larval samples, we extracted total DNA and RNA using both methods. Our data indicated that Method 1 harvested much lower concentration (*F*_*1, 28*_ = 16.2, *p* < 0.001) and yield than method 2 (*F*_*1, 28*_ = 15.0, *p* < 0.001, Figure 1).

**Figure 1:**
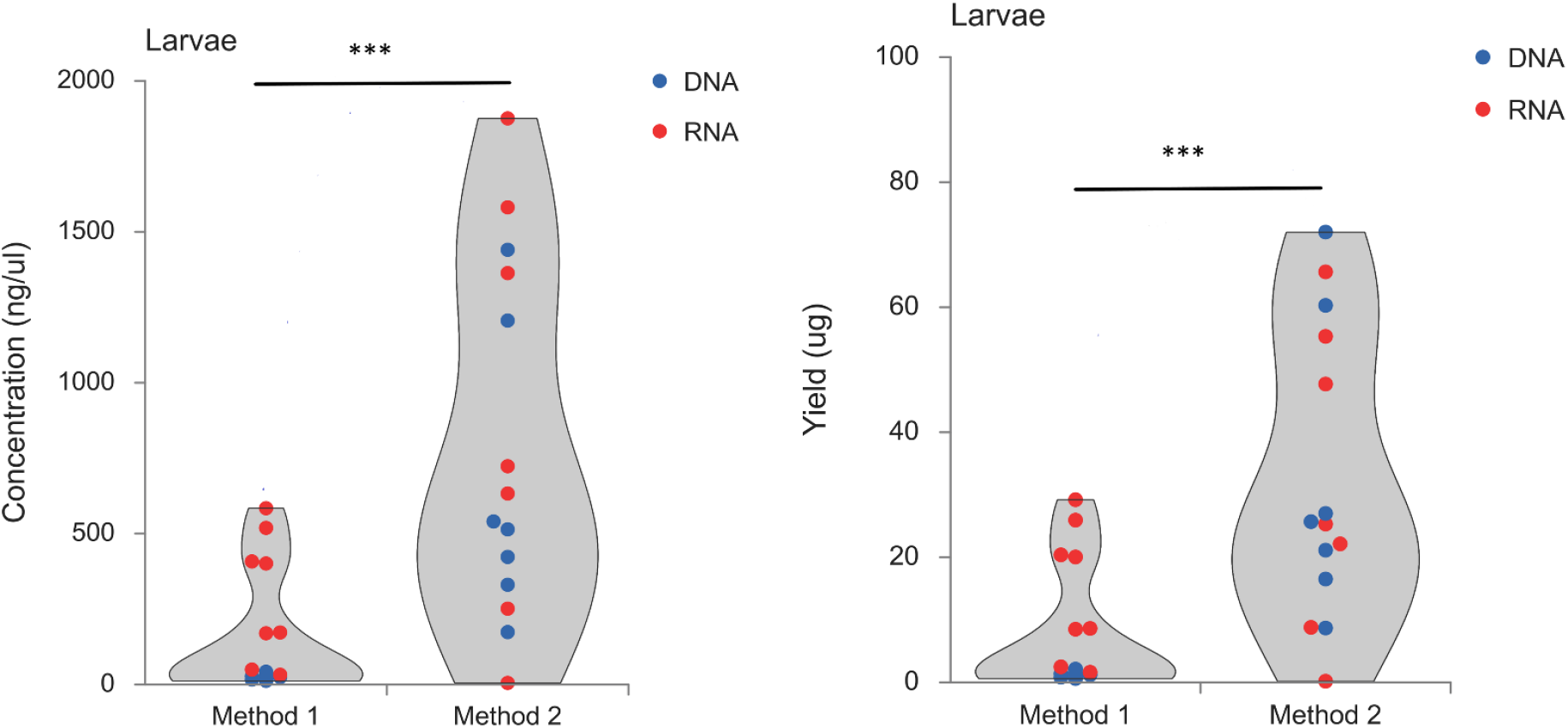
Comparison of concentrations (A) and yield (B) of total DNA and RNA per larvae sample.

In addition, we compared the pupal samples and extracted total DNA and RNA using both methods. For concentrations, there was a significant difference between the two methods (Figure 2). Concentrations of samples processed by Method 1 were much lower than Method 2 (*F*_*1, 26*_ =14.5, *p* < 0.001; Figure 2). We noticed a similar trend in total yield and DNA and RNA between the two methods. With a significant difference between the yield, samples from Method 1 showed a significant lower yield than Method 2 (*F*_*1, 26*_ = 12.1, *p* < 0.01; Figure 2).

**Figure 2:**
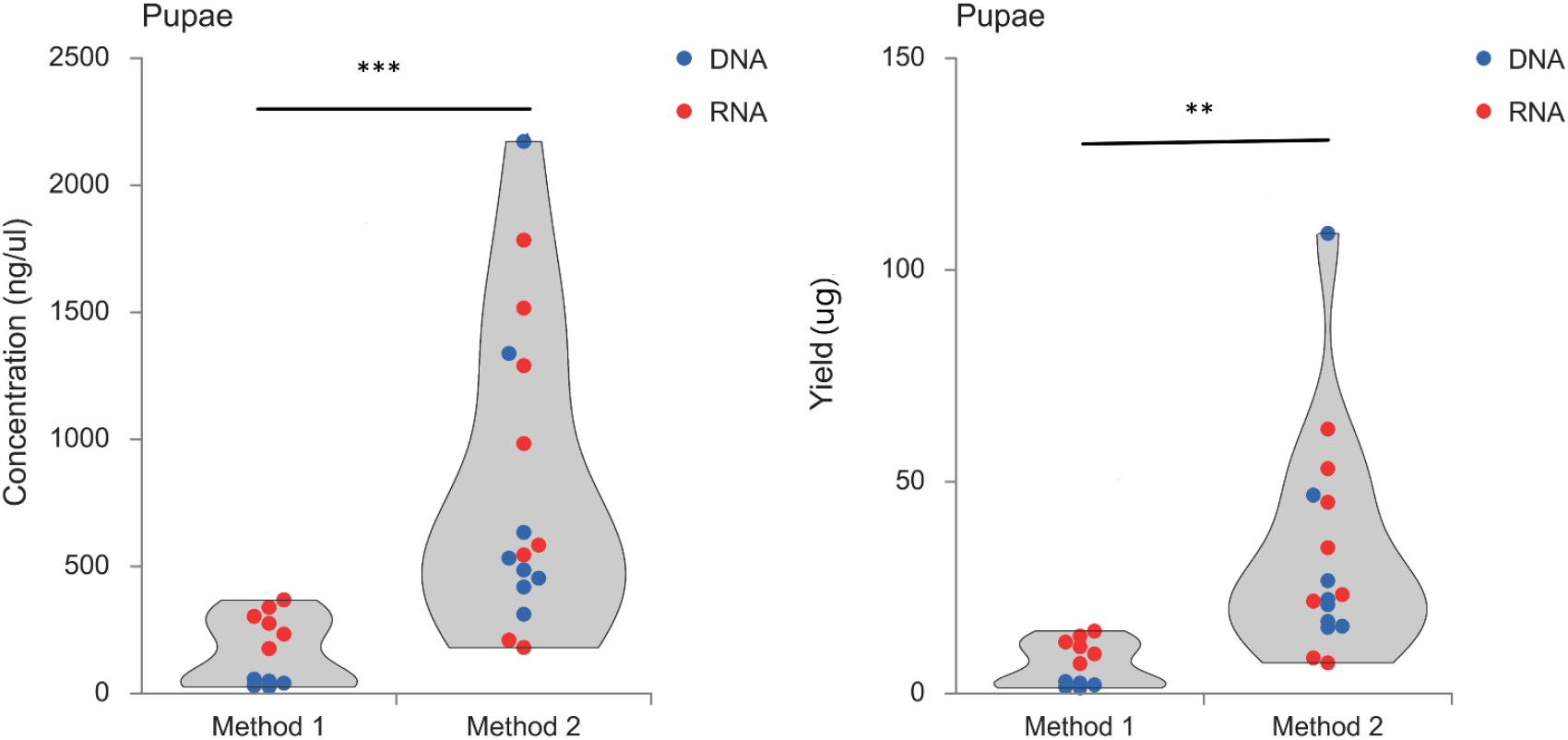
Comparison of concentrations (A) and yield (B) of total DNA and RNA per pupae sample.

To cover most developmental stages, we collected head tissues of adult worker bees to compare the concentration and yield. We continued to see the similar trend of significantly higher concentration (*F*_*1*,, *20*_ = 53.8, *p* < 0.001; Figure 3) and yield (*F*_*1, 20*_ = 47.1, *p* < 0.001; Figure 3) of samples processed in Method 2.

**Figure 3:**
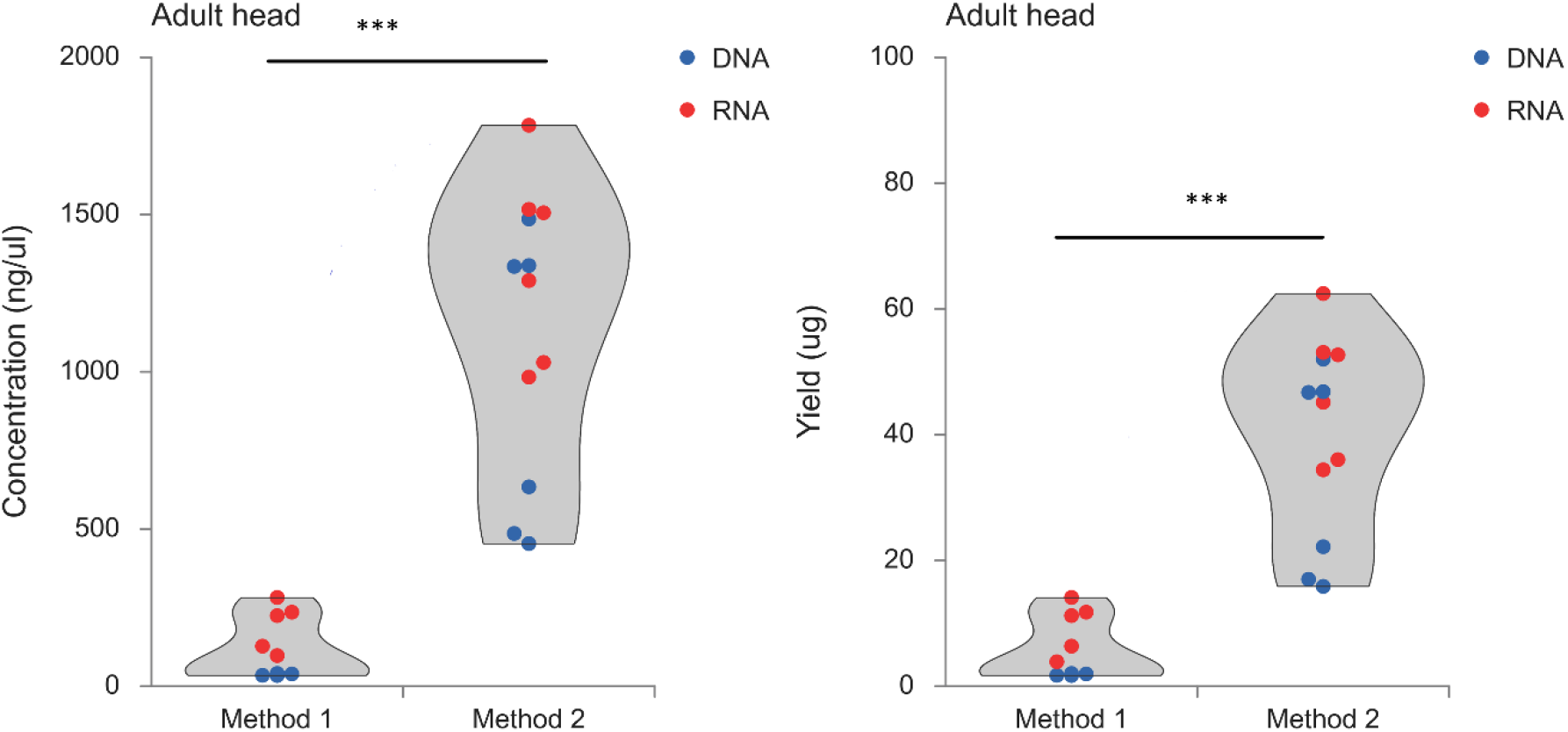
Comparison of concentrations (A) and yield (B) of total DNA and RNA per head of adult sample.

The quality of all samples was assessed by the ratio of A260/A280. All data exhibited a ratio between 1.7 and 2.2, which are acceptable for further experiments. The differences in quality between the two techniques are not significantly different for DNA (*F*_*1, 38*_ = 0.89, *p* = 0.35, Figure 4) and RNA (*F*_*1, 33*_ = 3.96, *p* = 0.055, Figure 4), respectively.

**Figure 4:**
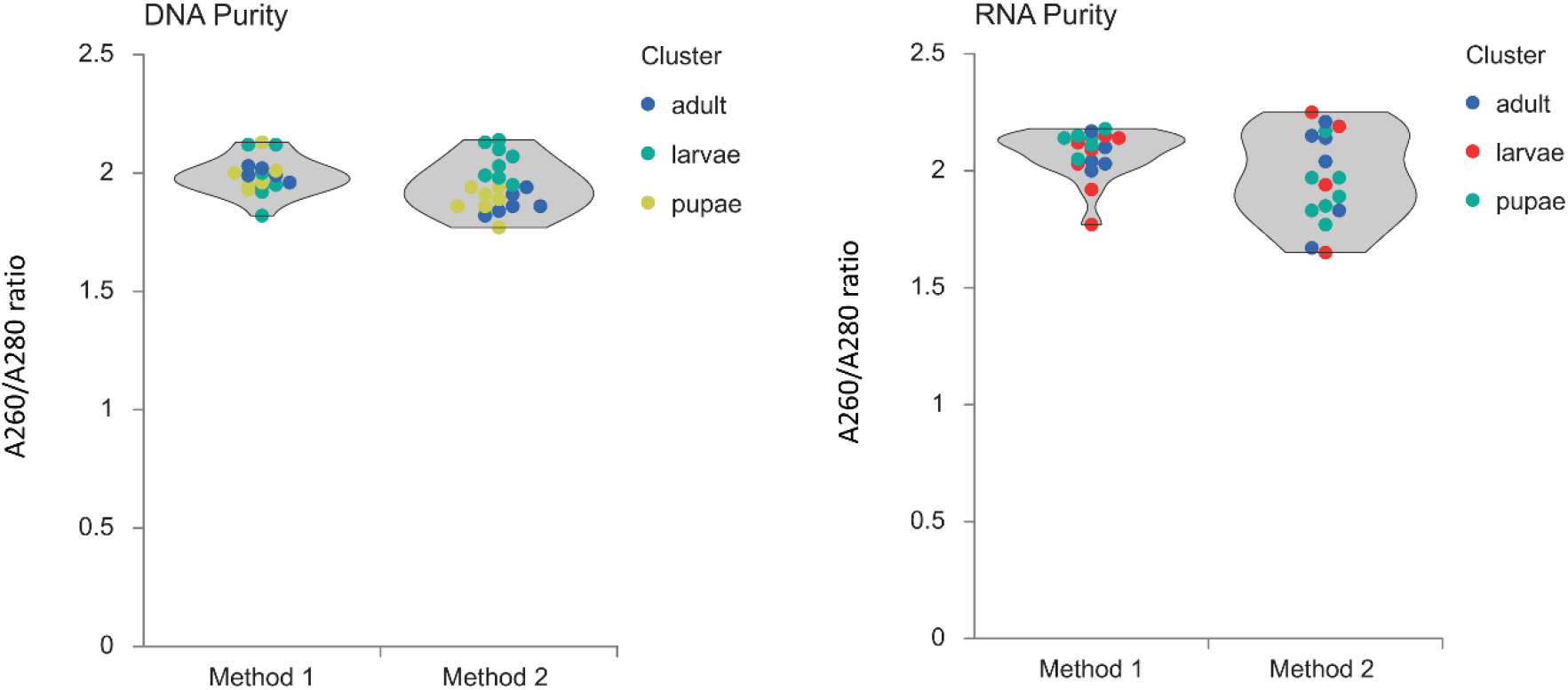
Comparison of purity in total DNA (A) and RNA (B) of all samples.

## 4. Discussion

Both methods enabled dual extraction of RNA and DNA from individual honey bees at the larval, pupal, and adult life stages. Comparing the DNA yield between the techniques and each life stage, Method 2 (following a more traditional procedure) produced significantly higher yields than Method 1 (using columns). This result extends to the RNA isolations; Method 2 showed significantly higher yields of RNA for the larval, pupal, and adult life stages.

There were some notable differences between the two dual extraction kits. For instance, the Method 1 had a faster turn-around time (40 minutes per sample, on average) and used a column-based technique. Method 2 took one hour per sample on average and uses a precipitation-based technique. Nonetheless, we demonstrate here that DNA and RNA can be dually extracted from individual honey bees. Although the yield is higher with Method 2, the faster extraction time and lower cost may make Method 1 more desirable for some applications.

If we consider the time for performing the experiments, it is usually faster to use the column based (Method 1) than the traditional alcohol precipitation (Method 2). However, Method 1 has two major drawbacks: (1) membrane spin columns cost more than traditional Method 2 of purification, and (2) membrane spin columns have specific abilities to select certain size of nucleic acid they can purify, excluding very large (>50 kb) or small nucleic acids (<100 bp) (Dowhan, 2012). Based on our comparison, we see a significant lower concentration and lower yield of DNA and RNA using Method 1, which are membrane spin columns. For future directions, comparison of the distribution of length between two methods are needed.

We did not study the integrity of the nucleic acid (e.g., length of the molecules), which may be an important criterion for genomic applications. In general, precipitation versus column methods are dependent on the desired aim of the study. When looking for the greatest yield, precipitation methods will be more effective than the column-based methods of extraction. Precipitation methods run into issues when discussing the overall purity of the extracted samples, which is when the column-based extractions are superior. The overall advantage of dual genomic extractions arises from the RNA providing an insight into the total gene expression of the individual at the moment of extraction, and the genome level expression of proteins within that individual as well. Further study into dual extractions amongst castes can provide greater insight into differences in gene expression as well as quantifying levels of mitochondrial DNA amongst castes. This type of information will provide more information into how social insects conduct interactions from a molecular level.

The future technology of genomic biology may focus on novel ways to sequence DNA and RNA simultaneously to gain a complete understanding of the genomic network and gene regulation in cells (Lee & Hwang, 2020). More social insects will be sequenced as the cost of sequencing decreases (http://antgenomics.dk/, http://i5k.github.io/) (Robinson et al., 2011). Our recent research of dual extraction of DNA and RNA reveals the importance of DNA methylation marks and how they may play an critical role in gene expression and alternative splicing (Li-Byarlay et al., 2020). In addition, previous studies showed investigations from both DNA and RNA sequencing levels are important to reveal novel molecular mechanisms for behavioral research of social insects (Foret et al., 2009, Herb et al., 2012, Standage et al., 2016, Galbraith et al., 2015).

In summary, we demonstrate that dual extraction is feasible for larval, pupal, and adult (head) life stages. The results also show that the Method 2 produced much higher yields of both DNA and RNA from the samples. It is possible to use individual honey bees from most stages of life to extract DNA and RNA with these protocols. Using separated samples of the same individuals (as opposed to pooled sample of different individuals) can increase precision and accuracy while decreasing inter-individual noise from data by eliminating genetic variations because of other intrinsic differences among individuals. Dual extraction of individuals and even separate tissues of individuals like the head is a viable option for honey bee research.

## Author Contributions

Conceptualization, HL-B and DRT; Methodology, RS, BG, and HL-B; Data Analysis, RS, BG, and HL-B; Resources, MS, DRT; Writing, Original Draft Preparation, BG, RS; Writing – Review & Editing, HL-B, DRT, MS; Funding Acquisition, HL-B and DRT.

## Acknowledgments

This research was supported by funding from the National Research Council via a Senior Research Associateship to HL-B, USDA-NIFA Evans Allen Program to HL-B and RS, NCSU Undergraduate Research Scholarship to BG.

